# Preferred hexoses influence long-term memory and induction of lactose catabolism by *Streptococcus mutans*

**DOI:** 10.1101/300749

**Authors:** Lin Zeng, Robert A. Burne

**Author notes:** Corresponding author: 1395 Center Drive, Gainesville, Florida 32607.

## Abstract

Bacteria prioritize sugar metabolism via carbohydrate catabolite repression, which regulates global gene expression to optimize the catabolism of preferred substrates. Here, we report an unusual long-term memory effect in certain *Streptococcus mutans* strains that alters adaptation to growth on lactose after prior exposure to glucose or fructose. In strain GS-5, cells that were first cultured on fructose then transferred to lactose displayed an exceptionally long lag (>11 h) and slower growth, compared to cells first cultured on glucose or cellobiose, which displayed a reduction in lag phase by as much as 10 h. Mutants lacking the cellobiose-PTS or phospho-β-glucosidase lost the accelerated growth on lactose associated with prior culturing on glucose. The memory effects of glucose or fructose on lactose catabolism were not as profound in strain UA159, but the lag phase was considerably shorter in mutants lacking the glucose-PTS EII^Man^. Interestingly, when *S*. *mutans* was cultivated on lactose, significant quantities of free glucose accumulated in the medium, with higher levels found in the cultures of strains lacking EII^Man^, glucokinase, or both. Free glucose was also detected in cultures that were utilizing cellobiose or trehalose, albeit at lower levels. Such release of hexoses by *S*. *mutans* is likely of biological significance as it was found that cells required small amounts of glucose or other preferred carbohydrates to initiate efficient growth on lactose. These findings suggest that *S*. *mutans* modulates the induction of lactose utilization based on its prior exposure to glucose or fructose, which can be liberated from common disaccharides.

**IMPORTANCE:** Understanding the molecular mechanisms employed by oral bacteria to control sugar metabolism is key to developing novel therapies for management of dental caries and other oral diseases. Lactose is a naturally occurring disaccharide that is abundant in dairy products and commonly ingested by humans. However, for the dental caries pathogen *Streptococcus mutans*, relatively little is known about the molecular mechanisms that regulate expression of genes required for lactose uptake and catabolism. Two peculiarities of lactose utilization by *S*. *mutans* are explored here: a) *S*. *mutans* excretes glucose that it cleaves from lactose and b) prior exposure to certain carbohydrates can result in a long-term inability to use lactose. The study begins to shed light on how *S*. *mutans* may bet-hedge to optimize its persistence and virulence in the human oral cavity.

## INTRODUCTION

Dental caries is a disease caused by metabolic activities of tooth biofilms, driven primarily by the fermentation by microorganisms of carbohydrates to organic acids that dissolve the tooth minerals. The development of caries usually requires ingestion of substantial amounts of dietary carbohydrates over an extended period of time. In developed nations, starches, sucrose, high-fructose corn syrups that contain a mixture of glucose and fructose, and lactose are among the most abundant carbohydrates in the human diet. Since feeding in humans is intermittent, the oral microbiota endures a “feast-or-famine” existence (1), wherein oral biofilms persist during fasting periods on nutrients provided in saliva and other oral secretions, end products generated by microorganisms, and microbially produced storage compounds that include intracellular glycogen-like polysaccharides (IPS) and structurally and compositionally diverse extracellular polysaccharides (EPS). The microbiome is generally believed to be carbohydrate-limited when the host is fasting. Most bacteria employ carbohydrate catabolite repression (CCR) to prioritize carbohydrate utilization when more than one metabolizable source is present, shifting to less-preferred carbohydrates after preferred sources have been exhausted. The primary effector of CCR in low G+C Gram-positive bacteria is usually catabolite control protein (CcpA), which differentially regulates a significant portion of the transcriptome in response to certain glycolytic intermediates (2), including fructose-1,6-bisphosphate (F-1,6-bP). The major etiological agent for human dental caries, *Streptococcus mutans*, is a Gram-positive, lactic acid-generating bacterium capable of metabolizing an array of carbohydrates, even at pH values as low as 3.8 (3). To persist and cause disease, *S*. *mutans* must compete with a variety of metabolically similar commensal streptococci and with other caries pathogens. With the recognition that dental caries is a polymicrobial infectious disease, there is a need to improve our understanding of the molecular mechanisms and the effects on bacterial ecology of the prioritization and optimization of carbohydrate utilization by the oral microbiome.

Significant progress has been made in understanding CCR in *S*. *mutans*, including the characterization of the CcpA-regulon and its role in regulating central carbon metabolism and virulence gene expression (4, 5). *S*. *mutans* is unusual, however, in that CcpA does not play a major role in CCR by directly regulating catabolic genes for non-preferred carbohydrates. Instead, CcpA-independent pathways are the primary routes for managing CCR in this human pathogen. First, a major role in CCR has been demonstrated for the AB domain of the primary glucose/mannose-specific PTS permease EII^Man^, encoded by *manL*, which regulates the expression of the fructanase operon (*fruAB*), a fructose-PTS operon (*levDEFG*), the cellobiose-PTS pathway (*celA* and *celRBCD*), and the lactose operon (*lacABCDFEG*) (6–9). A *manL* mutant of *S*. *mutans* UA159 showed significant defects in biofilm formation, acid production and competence development (10, 11). Similarly, a mutant defective in the sucrose-PTS permease (*scrA*) showed altered expression of the *fruAB* and *levDEFG* operons (12). HPr, and in particular the serine-phosphorylated form of the protein (HPr-Ser46-PO_4_) is also a potent effector of CCR for certain catabolic systems in *S*. *mutans*, as is the case for *B*. *subtilis* and other low G+C Gram-positives.

Due in large part to the widespread adoption of diets rich in refined carbohydrates, fructose and lactose have become increasingly abundant in human foods in recent decades. However, unlike for sucrose, the relative contribution of these and many other sugars to human dental caries and the overall impact on the oral microbiome is not well understood. Fructose is considered a preferred sugar for many oral bacteria and is catabolized by oral streptococci in a manner similar to glucose. Nearly 5% of the transcriptome of *S*. *mutans* is differentially expressed (>2-fold change in mRNA levels) in response to fructose, compared to cells growing on glucose (13), indicative of a major shift in the physiological state of cells when the two different hexoses are present. Notably, many genes encoding proteins needed for development of genetic competence and for stress tolerance showed enhanced expression in cells growing on fructose. A recent study probing the molecular mechanisms by which fructose affects gene expression in *S*. *mutans* revealed that FruR regulates expression of a primary fructose-PTS (*fruI*) and the glucose/mannose-PTS operon (*manLMN*) in concert with the CcpA protein (14). It was also reported that accumulation of phosphorylated fructose derivatives, particularly in a *fruK* (1-phosphofructokinase) mutant, adversely affected the ability of the organism to grow on multiple carbohydrates; and loss of FruK altered the expression of ~400 genes.

Lactose is a β1,4-linked disaccharide of glucose and galactose. Catabolism of lactose by bacteria usually requires cleavage of the disaccharide, either inside or outside of the cell, or following internalization and phosphorylation by the PTS. Lactose metabolism in *S*. *mutans* depends on the gene products of the *lac* operon, which encodes a lactose-specific PTS (*lacFE*) for transport and phosphorylation of lactose, a phospho-β-galactosidase (*lacG*) for cleavage of lactose-6-phosphate (Lac-6-P), and the tagatose pathway (*lacABCD*) for metabolism of the galactose-6-phosphate (Gal-6-P)(9, 15) that is released from Lac-6-P. Intracellular glucose is also generated that can be phosphorylated by a glucokinase (*glk*) and enter the glycolytic pathway. Expression of the *lac* genes requires activation by Gal-6-P, which derepresses the operon presumably by acting as an allosteric regulator of the LacI-type regulator LacR. The expression of the *lac* operon is also subject to negative regulation by the glucose-PTS and is repressed when sufficient levels of glucose are available in the environment (9). When both lactose and glucose are present, *S*. *mutans* preferentially internalizes and catabolizes glucose (16). After glucose is depleted, cells shift to the use of lactose for growth, with batch-cultured cells exhibiting a classic diauxic growth curve (16). The underlying mechanism for glucose-dependent repression appears to be inducer exclusion, whereby expression of the *lac* genes is inhibited due to a failure to transport lactose and cleave it to produce the apparent cognate inducer (Gal-6-P); however, other factors that may regulate repression and induction have not yet been elucidated (9). The tagatose pathway is the primary route for catabolism of galactose by *S*. *mutans*, but there are strain-specific differences that dictate whether galactose is transported by a galactose-specific PTS or the lactose-PTS, with EII^Man^ serving as a secondary lower-affinity transporter for galactose (9).

We posit that the ability of *S*. *mutans* to coordinate and optimize the utilization of preferred carbohydrates and non-preferred sources to compete with commensals and throughout the caries process is an essential virulence attribute. To better understand the molecular mechanisms used by *S*. *mutans* and related organisms, we examined the transition of *S*. *mutans* from preferred sugars to non-preferred sources using lactose as a model. The results from these experiments revealed an unusual memory and growth arrest phenomena, with glucose, fructose and the PTS regulating these behaviors.

## RESULTS AND DISCUSSION

### History of exposure to glucose or fructose affects adaptation to lactose

To investigate how *S*. *mutans* prioritizes carbohydrate metabolism, we performed a series of growth assays examining the transition from preferred sugars to lactose. Starting from an overnight culture prepared using a modified FMC medium that was constituted with a final concentration of 20 mM glucose or fructose, *S*. *mutans* cultures were diluted 1:10 into the same media, sub-cultured to mid-exponential phase, then diluted (1:20) into fresh FMC containing 10 mM lactose. Growth on lactose was monitored using a Bioscreen C for 48 h. *S*. *mutans* GS-5 and a number of other strains of *S*. *mutans* (10449, LM7, V403, and SMU44), did not display any appreciable growth for several hours (12-15 h for GS-5) after transfer to lactose when the cells were pre-cultured in FMC-fructose, but initiated growth within 2 h after transfer from FMC-glucose to FMC-lactose (Fig. 1A, Table 1, and Fig. S1A of supplementary materials). Most of these strains (except LM7) also exhibited slower growth and attained lower optical densities (Fig. S1A) following transfer from FMC-fructose to FMC-lactose, compared with cells transferred from FMC-glucose. Similar behaviors were observed under conditions when cells were transferred to a mixture of 5 mM lactose and 5 mM glucose or fructose (Fig. S2), where significantly slower growth was noted from cells pre-cultivated on fructose, when they were transferred onto FMC containing a mixture of glucose and lactose. These results suggest that growth on fructose induced a form of long-term “memory” that prevented cells from initiating growth on lactose. This effect is not a simple manifestation of CCR, since cells transitioned in a much more typical fashion from glucose to lactose, and the effects of pre-cultivation in fructose persisted even when glucose was used alone, in place of lactose, or added to the FMC-lactose (data not shown and Fig. S2).

**Figure 1.**
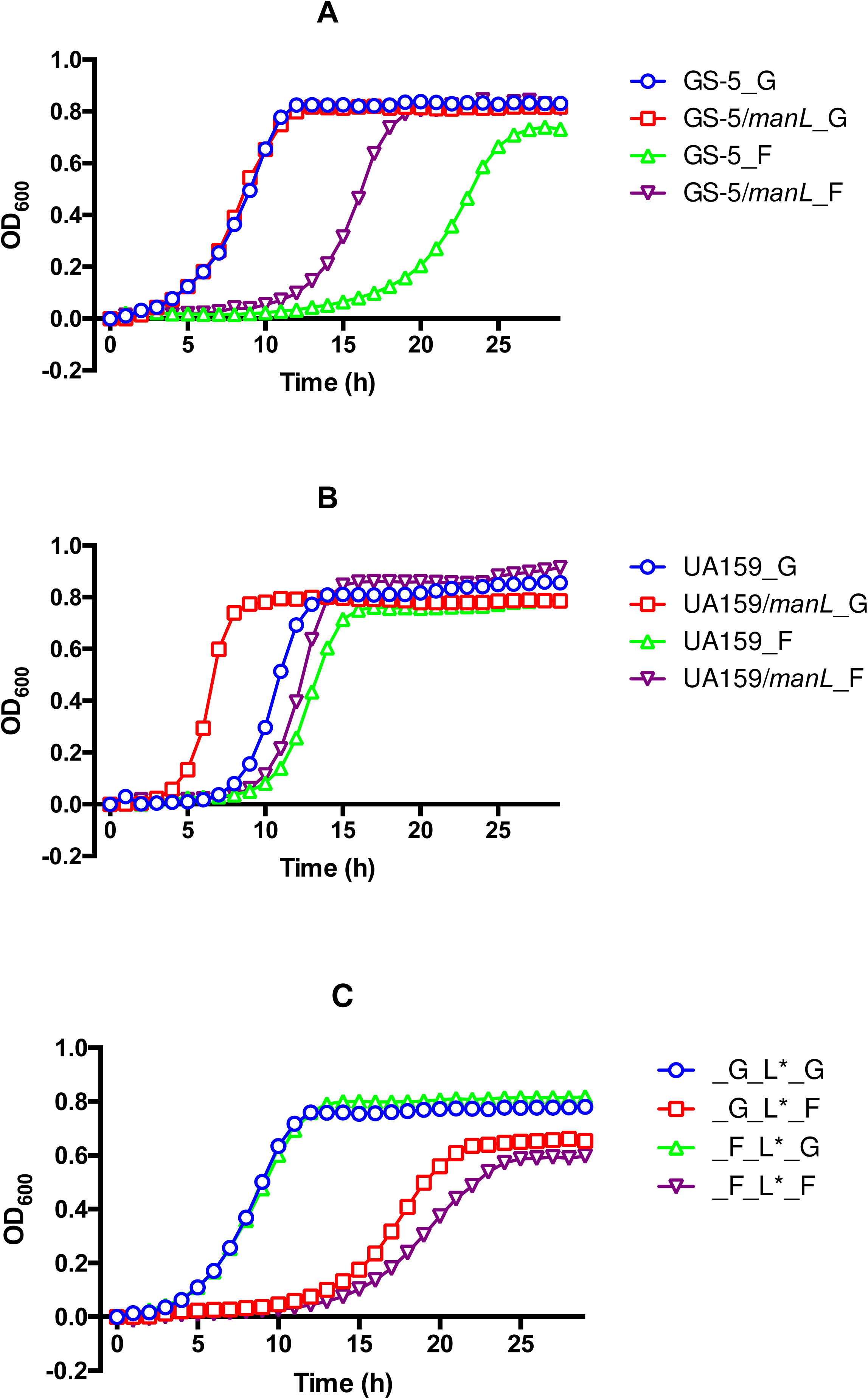
Growth of *S*. *mutans* on lactose after transferring from glucose or fructose. Wild-type strains GS-5 (A) and UA159 (B) and their otherwise-isogenic mutants deficient in glucose-PTS permease (*manL*) were first cultivated in FMC containing 20 mM glucose (designated as _G) or fructose (_F) till exponential phase, then diluted at 1:20 ratio into fresh FMC constituted with 10 mM lactose for growth monitoring on a Bioscreen C system. (C) The final cultures from (A) were spread on BHI agar plates to form individual colonies, designated as _G_L^∗^ and _F_L^∗^, which were cultivated again on glucose (_G) or fructose (_F) and diluted into FMC containing lactose.

**Table 1.** Growth characteristics of *S*. *mutans* wild-type strains, their isogenic mutants, and *S*. *gordonii* in FMC medium containing 10 mM lactose^a^.

| Strains |  | Growth on lactose |  |  |  |
| --- | --- | --- | --- | --- | --- |
| | | Carbohydrates for pre-culture | $T_d$ (min) | OD<sub>600</sub> | $T_{lag}$ (h) |
| UA159 | wild type | Glucose | 66.0 ± 2.1 | 0.87 | 5.5 |
|  |  | Fructose | 74.9 ± 5.7 | 0.80 | 7.5 |
|  |  | Cellobiose | 96.1 ± 6.7 | 0.82 | 3.0 |
|  |  | Galactose | 56.2 ± 5.1 | 0.87 | 1.5 |
|  |  | Lactose | 52.7 ± 4.4 | 0.94 | 1.0 |
|  | JAM1 (*manL*) | Glucose | 57.5 ± 1.2 | 0.81 | 2.0 |
|  |  | Fructose | 70.1 ± 3.4 | 0.91 | 7.0 |
|  | *manL celR* | Glucose | 66.3 ± 3.9 | 0.74 | 6.5 |
|  |  | Fructose | 69.0 ± 2.6 | 0.81 | 6.0 |
|  | *relA::Km* | Glucose | 75.2 ± 5.9 | 0.74 | 10.5 |
|  | *relA*<sup>385</sup> | Glucose | 80.6 ± 3.6 | 0.77 | 11.0 |
|  | *relP* | Glucose | 63.2 ± 3.0 | 1.00 | 5.0 |
|  | *relQ* | Glucose | 65.5 ± 0.9 | 0.98 | 5.5 |
|  | *relAPQ* | Glucose | 69.7 ± 0.2 | 0.99 | 8.0 |
| GS-5 | wild type | Glucose | 119.5 ± 1.3 | 0.84 | 1.5 |
|  |  | Fructose | 150.1 ± 1.1 | 0.73 | 11.5 |
|  |  | Cellobiose | 121.8 ± 9.6 | 0.70 | 2.5 |
|  |  | Galactose | 160.7 ± 4.3 | 0.58 | 2.0 |
|  |  | Lactose | 167.5 ± 8.0 | 0.66 | 2.0 |
|  | *manL* | Glucose | 108.0 ± 0.03 | 0.83 | 2.0 |
|  |  | Fructose | 105.0 ± 4.9 | 0.84 | 5.5 |
|  | *celR* | Glucose | 68.1 ± 5.6 | 0.90 | 6.0 |
|  |  | Fructose | 68.0 ± 3.7 | 0.93 | 6.0 |
|  | *celA* | Glucose | 109.1 ± 6.1 | 1.13 | 2.5 |
|  |  | Fructose | 287.5 ± 14.6 | 0.48 | 9.5 |
|  | *celB* | Glucose | 183.5 ± 14.7 | 0.57 | 9.5 |
|  |  | Fructose | 180.2 ± 1.0 | 0.58 | 9.0 |
| *S. gordonii* DL1 |  | Glucose | 49.6 ± 3.0 | 0.82 | 0.5 |
|  |  | Fructose | 51.8 ± 3.4 | 0.82 | 0.5 |
|  |  | Cellobiose | 48.4 ± 3.5 | 0.93 | 0.5 |
|  |  | Lactose | 50.1 ± 4.0 | 0.91 | 0.5 |

<sup>a</sup> Each culture was started by 1:20 dilution into FMC-lactose of exponentially growing cultures prepared with FMC containing the specified carbohydrate. Each doubling time ( $T_d$ in min, average ± standard deviation) was calculated as $T_d = (T_2 - T_1) \times \text{LOG}(2) / \text{LOG}(OD_2 / OD_1)$ using data from exponential phase ( $OD_{600} = 0.2 \sim 0.4$ ). Maximal optical density ( $OD_{600}$ ) and the length of the lag phase ( $T_{lag}$ in hour) reported were the averages of results from at least three biological replicates. The end of the lag phase was defined as the point at which $OD_{600}$ reached 0.01.

The results above contrasted with what was observed with the commonly utilized laboratory strain UA159 and certain other isolates of *S*. *mutans* (UA101, SMU33, SMU21, and OMZ175) (Fig. 1B, Table 1, and Fig. S1B). In particular, the times required to initiate growth on lactose following transfer from fructose were much shorter than for the strains described in Figure S1A, with no long-term memory being evident. However, the time it took for glucose-grown cells to initiate growth on lactose increased for these strains, except for OMZ175.

Glucose is internalized primarily by the glucose/mannose-PTS transporter EII^Man^, so the growth behaviors of EII^Man^ mutants, generated by deleting *manL* or *manLMN*, in the UA159 and GS-5 genetic backgrounds were also evaluated. The results (Fig. 1B and Table 1) showed that, when compared with the wild-type strains, the EII^Man^ mutant of UA159 (JAM1, deficient in *manL*) showed a more rapid resumption of growth following transfer from FMC-glucose to FMC-lactose, with a reduction in lag phase by as much as 3.5 h. However, JAM1 transitioned from fructose to lactose with a lag time similar to UA159. Interestingly, the EII^Man^ mutant of GS-5 adapted much faster to growth on lactose from fructose (~ 6 h reduction in lag), but not from glucose to lactose, when compared with the parental strain (Fig. 1A). These observations provide further evidence that metabolism of glucose or fructose can influence the imprinting of “memory” in the organisms that prevents the population from metabolizing lactose. It is interesting that strain-specific behaviors are observed in terms of the role of EII^Man^ in the behaviors of interest. We posit that growth on glucose results in the expression of genes that generate metabolic products that can promote lactose utilization by the bacterium, whereas the changes in gene expression and intermediates resulting from growth on fructose appear to significantly inhibit the expression or activity of gene products required for efficient transition to growth on lactose.

Experiments were also performed to exclude the possibility that cells that eventually grew in lactose after a prolonged lag were genetic variants, e.g. mutants that aberrantly regulated expression of *lac* genes. In particular, *S*. *mutans* GS-5 was cultivated overnight on FMC containing 20 mM glucose or fructose, sub-cultured on FMC with 10 mM lactose to late exponential phase. Cultures were then spread on BHI agar plates to form individual colonies, designated as _G_L^∗^ and _F_L^∗^, which were subsequently monitored for their growth behaviors during the transition from glucose or fructose to lactose. Consistent with the fact that these newly formed colonies were not genetic variants, the isolates behaved like the parental strain and (Fig. 1C) again showed that glucose-grown cells could transition onto lactose more rapidly than those cultured in fructose, as described above (Fig. 1A).

### Memory effects from other carbohydrates

To better understand why glucose and fructose have different effects on lactose metabolism, we tested how other carbohydrates influence the transition to lactose, first by cultivating *S*. *mutans* strains to exponential phase in the sugars of interest, then diluting the cultures into fresh FMC-lactose and monitoring growth. Cellobiose was tested out of the consideration that EII^Cel^ can function as an inducible glucose-PTS in *S*. *mutans* (7), and galactose was tested as it is primarily catabolized via the tagatose pathway, which is encoded in the *lac* operon and is the primary route for utilization of the galactose moiety of lactose. Lactose was also tested with the assumption that it should ensure optimal induction of the *lac* operon and that growth would initiate quickly after cells were transferred from FMC-lactose to the same medium. *S*. *mutans* UA159 cultivated in lactose or galactose rapidly transitioned onto lactose, with lactose-grown cells doing so significantly faster than glucose-grown cells and somewhat more rapidly than cells pre-cultured in galactose (Table 1 and Fig. S3A). Interestingly, pre-culturing on cellobiose also significantly enhanced the transition to lactose, more so than glucose, but not as much as lactose or galactose (Table 1).

Surprisingly, pre-cultivation in cellobiose had the greatest impact on the ability of strain GS-5 to transition to lactose, with cultures showing a shorter lag and faster growth rate on lactose (Table 1 and Fig. S3B) than cells pre-cultured on lactose or galactose. Again, pre-cultivation with lactose or galactose had nearly identical effects, and the transition in this case was similar to cells that were first cultured in glucose. Therefore, similar to exposure to glucose, pre-cultivation of *S*. *mutans* on cellobiose also significantly enhanced its capacity to transition to lactose, as compared to cells pre-cultured with fructose. Notably, the PTS permeases for cellobiose (EII^Cel^) and lactose (EII^Lac^) share significant sequence similarities and in some organisms each permease can transport both cellobiose and lactose; as noted for *Lactococcus lactis* IL1403, where EII^Cel^ and a phospho-β-glucosidase (BglS) cooperate to internalize then cleave lactose-6-P, respectively (17, 18). It is plausible, then, that induction of expression of the cellobiose-PTS by glucose or cellobiose leads to enhanced growth on lactose because EII^Cel^ is already induced and can rapidly internalize lactose, thereby providing inducing intermediates that trigger *lac* gene expression.

### Commensal oral streptococci do not display memory

As a constituent of the oral microbiota, *S*. *mutans* must compete against metabolically similar commensal bacteria for carbohydrates. To begin to investigate how memory in utilization of carbohydrates by *S*. *mutans* may influence oral microbial ecology, we performed the same experiments using a common oral commensal, *Streptococcus gordonii* DL1. Interestingly, DL1 grew efficiently on lactose when transitioning from glucose or fructose, with little difference in lag time, growth rate or final yield (Table 1 and Fig. S4). Similar results were obtained when *Streptococcus intermedius, Streptococcus sanguinis*, or *Streptococcus salivarius* were evaluated under similar conditions (Fig. S4), raising the question of whether the memory behavior may be unique to *Streptococcus mutans* and possibly to cariogenic streptococci. Interestingly, for another *L*. *lactis* strain ATCC 11454 that is similar to IL1403, pre-cultivation on glucose (as compared to fructose) appeared to result in substantially improved growth on lactose (Fig. S4). Of note, a recent study with *Streptococcus pneumoniae* demonstrated that the glucose-PTS encoded by *manLMN* is required for the catabolism of a number of secondary carbohydrates, potentially by serving as a multi-substrate transporter and facilitating the induction of necessary catabolic genes (27). However, a comparable role for ManLMN of *S*. *mutans* appears unlikely since mutations in the *man* operon results in better growth on non-preferred carbohydrates.

### Defects in cellobiose metabolism alter glucose-induced memory

In *S*. *mutans*, the expression of the cellobiose pathway, which includes two transcripts encoding *celA* and *celBRCXD*, is subject to EII^Man^-mediated repression when glucose is present. The operon is induced in the presence of cellobiose alone, or it can be induced with glucose in a strain lacking a functional EII^Man^. In the absence of EII^Man^, EII^Cel^ becomes the primary glucose transporter (7). Induction of the *cel* operon is controlled by two phosphorylation events targeting the transcriptional activator, CelR. Phosphorylation of CelR by EII^Cel^ at His284 and His391 leads to inhibition of CelR activity, whereas phosphorylation by Enzyme I of the PTS at His226, His332 and His576 is required for CelR-dependent activation of the *cel* operon (7). When glucose is transported by the primary permease EII^Man^, EI preferentially phosphorylates EII^Man^ to optimize glucose utilization, which prevents induction of *cel* genes because CelR is not phosphorylated at His226, His332 and His576 by EI. However, when EII^Cel^ is engaged in transport of glucose, CelR is no longer phosphorylated at His284 and His391 by EII^Cel^ and is able to activate *cel* gene expression (7).

We first investigated the effects of a *celR* deletion on memory. As shown in Fig. 2A, a *celR* mutant of GS-5 no longer displayed enhanced growth on lactose when transitioning from glucose, and the mutant retained the slow transition from fructose to lactose noted with the parental strain. The same was true when a *celR* deletion was introduced into an EII^Man^ mutant of UA159 (UA159/*manL/celR*, Fig. 2B). However, the *celR* mutant of GS-5 (but not that of UA159) displayed a significant reduction in doubling time during exponential phase in comparison with the wild type (Table 1). Collectively, these results affirmed the notion that the glucose-associated impact on the transition to lactose occurs through the cellobiose pathway. To further differentiate the contribution to lactose metabolism by the EII^Cel^ (*celBCD*) and the phospho-β-glucosidase (*celA*), mutants deficient in *celB* or *celA* were constructed in strain GS-5. Growth tests using these mutants indicated that loss of *celA* resulted in slightly longer transition to, but significant enhancement in yield on, lactose (Fig. 2C). Loss of EII^Cel^ (*celB*) drastically increased the length of the lag (by ~8 h) required for GS-5 to transition from glucose onto lactose (Fig. 2C), suggesting that EII^Cel^ plays a more prominent role than does phospho-β-glucosidase (CelA) in lactose metabolism.

**Figure 2.**
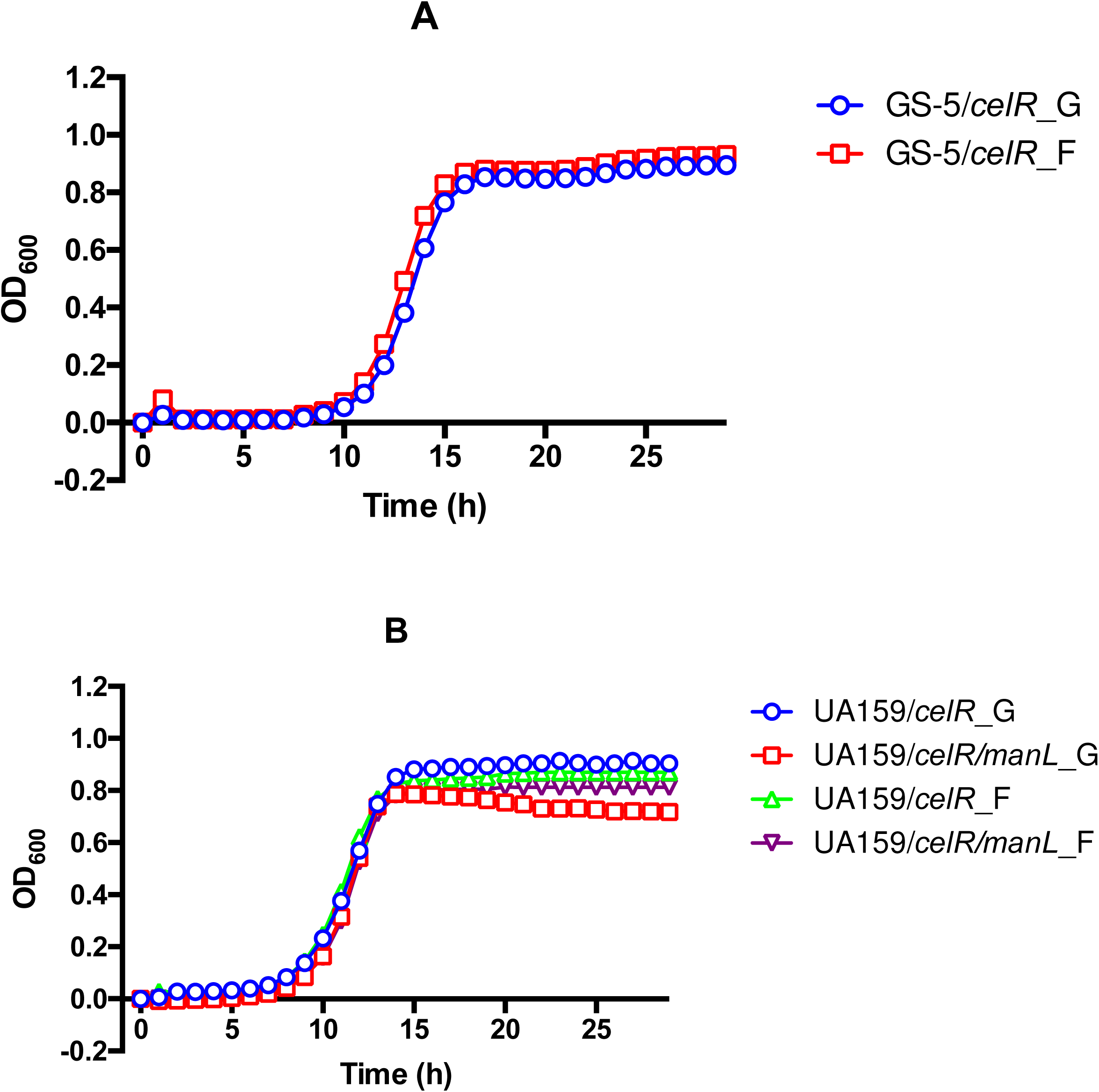
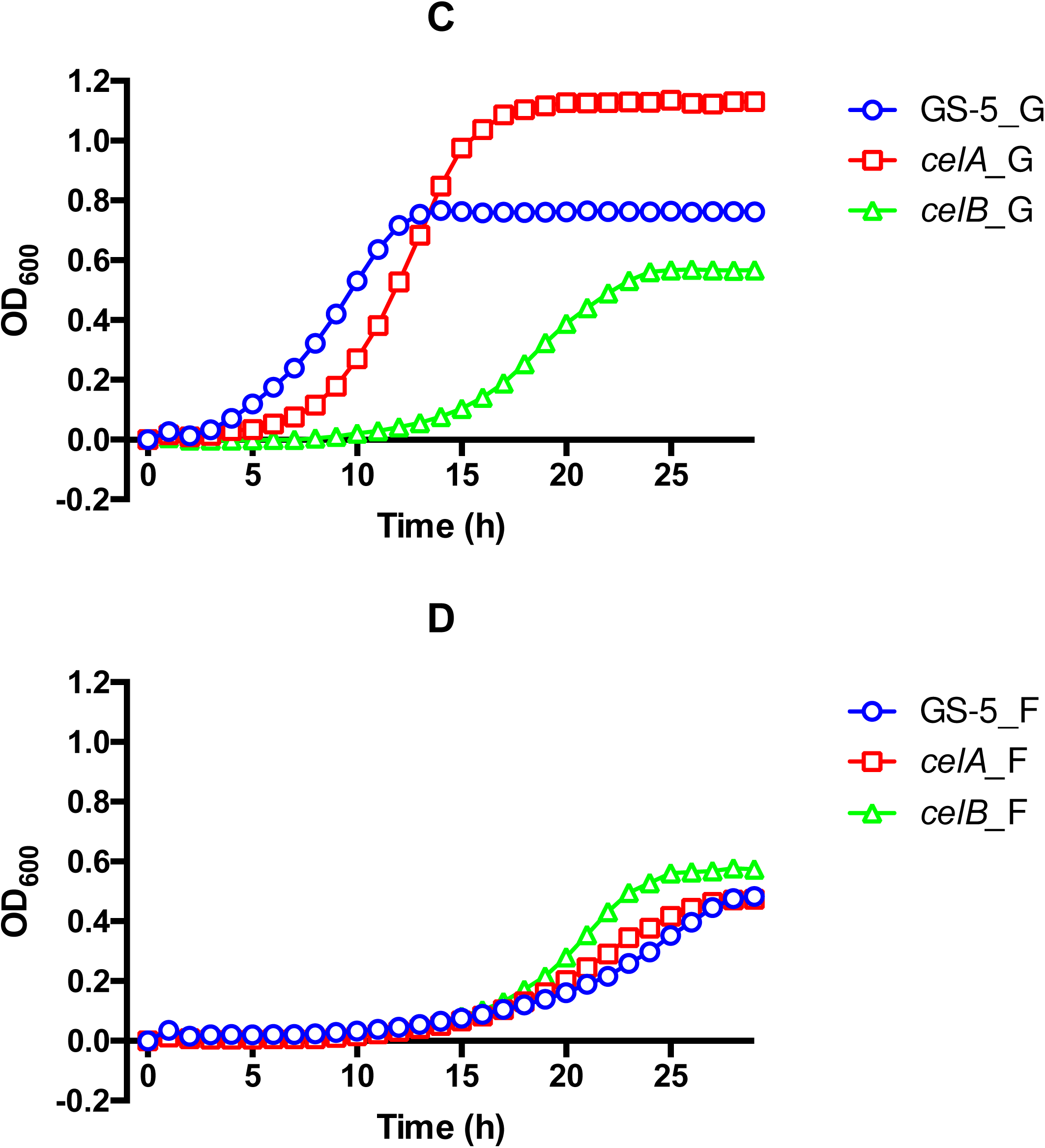
Disruption of the cellobiose pathway abolishes glucose memory. All strains were first cultivated in FMC containing 20 mM glucose (designated as _G) or fructose (_F) to mid-exponential phase (OD_600_ = 0.5), then diluted 1:20 into FMC containing 10 mM lactose for growth monitoring. (A) A *celR* mutant of GS-5. (B) Mutants derived from UA159 that are deficient in *celR*, or both *celR* and *manL*. (C) and (D), wild-type strain GS-5 and its otherwise-isogenic mutants deficient in phospho-β-glucosidase (*celA*) or EIIB of the cellobiose-PTS (*celB*) were grown in FMC with glucose (C) or fructose (D) before inoculation.

When fully induced, the cellobiose pathway of *L*. *lactis* alone is sufficient for growth on medium containing lactose as the carbohydrate source (18). It was posited that lactose is internalized by the cellobiose-PTS and phosphorylated on the glucose moiety, instead of on the galactose moiety; the latter being the case for lactose internalized by the lactose-PTS. Subsequent hydrolysis of the phosphorylated lactose moiety by BglS yields glucose-6-phosphate (G-6-P) and galactose, with galactose being catabolized by the Leloir pathway. To determine if a similar pathway exists in *S*. *mutans*, mutants deficient in lactose catabolic enzymes were constructed in the UA159 and GS-5 genetic backgrounds. Interestingly, the isogenic mutants of UA159 defective in EII^Lac^ (*lacE*) or the phospho-β-galactosidase (*lacG*) failed to grow on lactose, even when pre-cultured with cellobiose (Fig. S5). The same was true for the mutants derived from GS-5. Nevertheless, a previous analysis of lactose and galactose metabolism by *S*. *mutans* UA159 indicated a substantial induction (~6-fold increase in mRNA levels) of the Leloir pathway genes in cells growing on lactose compared to glucose, and such induction generally requires that the cells internalize galactose (9). Taken together, these findings suggest that the cellobiose pathway in *S*. *mutans* is not an independent, secondary catabolic pathway for lactose, but that the cellobiose PTS has a significant role in regulation of the catabolism of lactose; apparently by transporting and concomitantly phosphorylating lactose.

### Strain-specific effects of EII^Man^ in regulating *lac* gene expression

Growth phenotypes of the EII^Man^ mutants so far have shown alterations in the transition onto lactose under certain conditions (Fig. 1), which could be caused by relaxed repression of the *lac* operon, and in particular of the genes encoding the enzymes of the tagatose pathway (*lacABCD*). To test this theory, expression of the *lac* operon was investigated using a P*lacA::gfp* promoter fusion established in the wild-type and EII^Man^-deficient genetic backgrounds. Of note, we identified that there was a potentially significant difference in the sequence of the regions upstream of the *lacA* gene in strains UA159 and GS-5. In particular, two contiguous (T)_7_G motifs are present 193-nt upstream of *lacA* in GS-5, but only one is found in UA159 (Fig. S6), and no divergence is found in the rest of the *lacA* sequence. We therefore created two separate versions of *gfp*-promoter fusions, one containing two (T)_7_G motifs and the other containing one, and assessed GFP expression from both constructs in their cognate genetic backgrounds. Strains were first cultivated to mid-exponential phase in FMC-glucose, and then diluted into FMC containing only lactose or a mixture of equal concentrations of lactose and glucose. Growth and GFP levels were then monitored in real-time during in a BioTek Synergy plate reader (Fig. 3).

**Figure 3.**
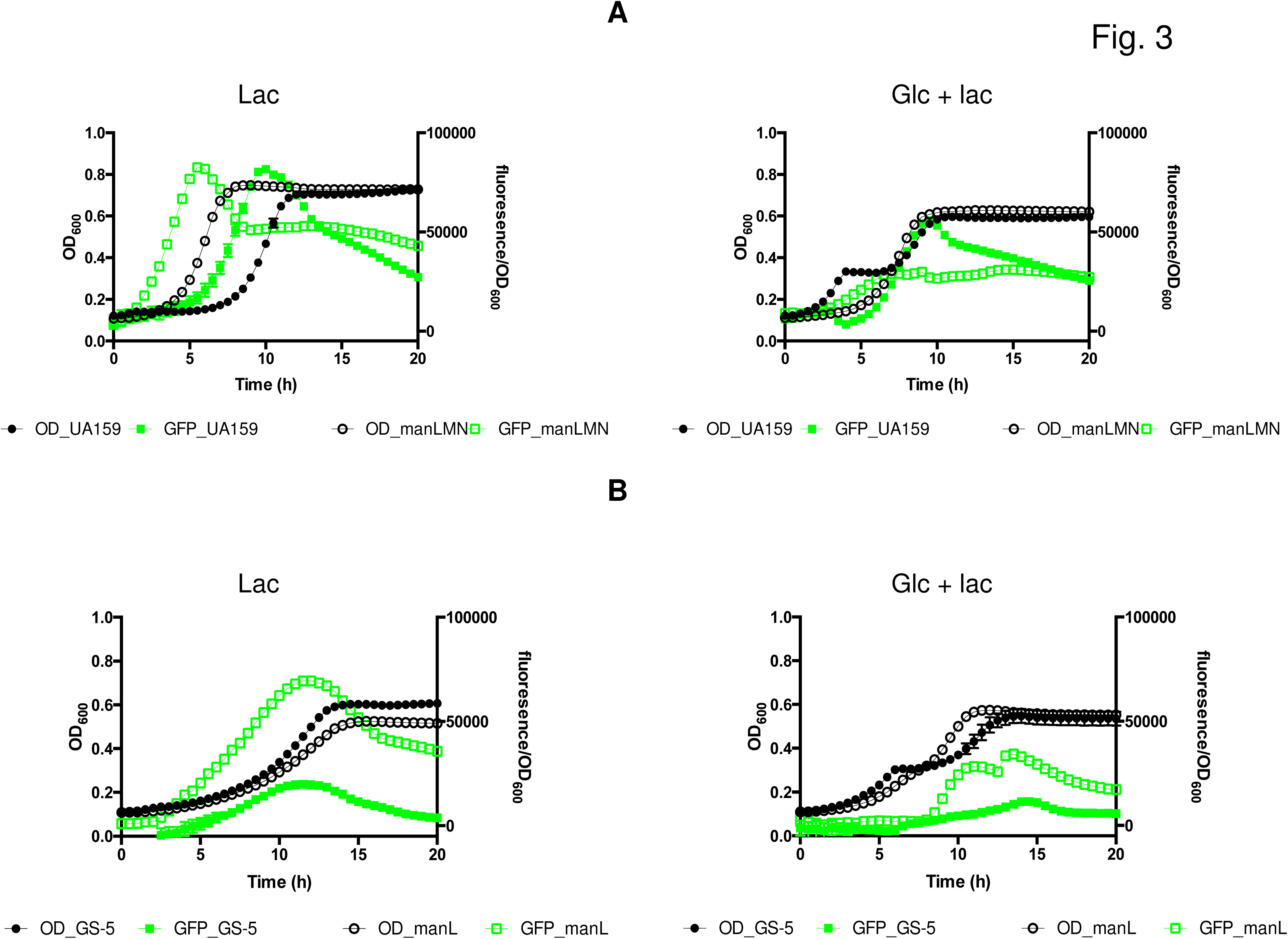
Growth curves (OD_600_, black) and expression of P*lacA::gfp* fusion (relative fluorescence, green) by strains in the UA159 (A) and GS-5 (B) backgrounds. Wild types (filled circle) and the mutants deficient in EII^Man^ (open circle) were cultivated to exponential phase in FMC containing 20 mM glucose, and then diluted into FMC containing 10 mM lactose (Lac) or 5 mM each of glucose and lactose (Glc + Lac). OD_600_ and GFP levels were monitored using a BioTek Synergy 2 plate reader maintained at 37°C. See methods section for information on normalization of the data.

Compared to what was seen in the wild-type UA159 genetic background, the EII^Man^ mutant (*manLMN;* and similarly *manL*, data not shown) showed a more rapid induction of the *lac* operon (by ~5 h), although the maximal levels of expression of GFP measured were similar in both strains (Fig. 3A). When the cells were monitored in a medium containing 5 mM glucose and 5 mM lactose, the wild type expectedly showed delayed *lac* expression until the onset of diauxie. In contrast, the EII^Man^ mutants presented no diauxie and showed constitutive, albeit significantly lower expression of the P*lacA::gfp* fusion, which started to increase early during growth but quickly plateaued in the mid-exponential phase of growth (OD_600_ ≈ 0.3); reaching levels significantly lower than that in the wild-type background (Fig. 3A). While the early induction of the *lac* operon in the EII^Man^-negative background can be viewed as a result of loss of inducer exclusion, the premature halt of such induction cannot be attributed simply to a loss of CCR, as the impact would have been the opposite.

Different from UA159, a GS-5 variant carrying its cognate P*lacA::gfp* promoter fusion, the one with two (T)_7_G motifs, produced generally lower GFP signals under the same conditions in comparison with UA159 carrying its cognate P*lacA::gfp* fusion. Importantly, the EII^Man^ mutant (*manL*) of GS-5 produced markedly higher expression of P*lacA::gfp* fusion, when compared with the wild type (Fig. 3B), while growing on lactose alone or in medium containing both lactose and glucose. However, as indicated earlier (Fig. 1A), this mutant did not show a shorter lag nor earlier *lac* gene induction during the transition from glucose to lactose. This phenotype will be discussed further below.

### Release of free glucose during lactose metabolism and its impact on gene regulation

In *S*. *mutans*, lactose, cellobiose and maltose are internalized by the PTS and the resultant phospho-disaccharides are cleaved to generate phosphorylated hexose and free glucose (7, 9, 19). The glucose thus produced can enter glycolysis following phosphorylation by a glucokinase (*glk*). Recent studies of the metabolism of maltose and maltooligosaccharides by *S*. *mutans* suggest that significant amounts of glucose are released into the environment and re-internalized by the PTS (20). We reported a similar release of free hexoses (fructose) when sucrose is catabolized through the sucrose-specific PTS by *S*. *mutans* (12, 13). Here, we show that when cultivated on lactose alone, free glucose can be detected in the supernates of the *manL* mutants in concentrations ranging from 0.5 to 1.5 mM/OD_600_. However, high levels of free glucose were not detectable in the supernatant fluid of the parental strain UA159, presumably because of re-internalization of the glucose by EII^Man^ in the wild type (Fig. 4). Notably, the concentrations of glucose that accumulated in the supernates were higher in strains carrying a deletion of *glk*, both in the wild-type and in the *manL* mutant genetic backgrounds (Fig. 4). Importantly, the presence of free glucose in the extracellular environment could affect the transcription of the cellobiose and *lac* operons. Specifically for UA159, small amounts of glucose transported by EII^Cel^ could induce a basal level expression of the cellobiose pathway without triggering CCR, and the *cel* gene products could then internalize lactose, similar to what is observed in *L*. *lactis*, but without generating Gal-6-P to induce the *lac* operon. In the absence of an intact EII^Man^, a situation that forces glucose internalization through EII^Cel^, extracellular glucose would further enhance the activities of the cellobiose pathway, significantly diverting the influx of lactose through the Cel system and resulting in down-regulation of the *lac* operon. This hypothesis was supported by the reduced expression of the P*lacA::gfp* fusion in the EII^Man^ mutant compared to cells with the wild-type genetic background, when growing on a mixture of both glucose and lactose (Fig. 3A).

**Figure 4.**
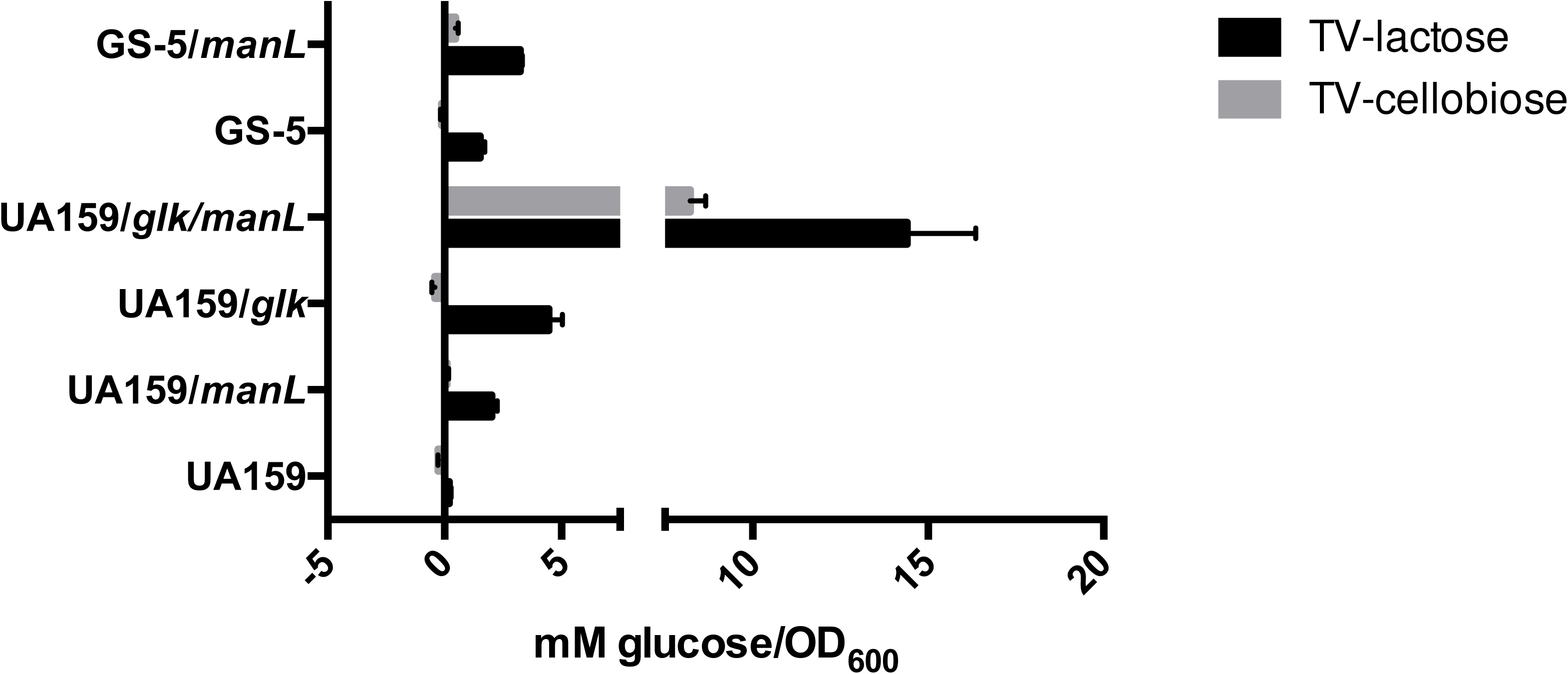
Detection of free glucose in the cultures of *S*. *mutans* growing on lactose or cellobiose. Wild-type strains UA159 and GS-5, as well as their isogenic mutants deficient in the glucose-PTS (*manL*) or glucokinase (*glK*), or both, were cultured in FMC containing 10 mM lactose or cellobiose to midexponential phase, then the supernates of the cultures were obtained after centrifugation and filtration. Levels of glucose in these supernates were measured using glucose assay kits. Each bar represents the average of three independent experiments that were each repeated at least twice, with error bars denoting the standard deviation.

To study the activities of the cellobiose operon, a P*celA::cat* promoter fusion was tested in the *manL* mutant. In agreement with our hypothesis, the P*celA::cat* fusion in the *manL* mutant of UA159 presented significantly increased activity compared to the wild type when growing on lactose, but especially when growing on a mixture of glucose and lactose (Fig. 5A). Furthermore, we have observed an apparent preference by both UA159 and GS-5, when growing on a mixture of equal amounts of glucose and lactose, to express the glucose and cellobiose PTS operons, rather than the lactose-PTS (Fig. 5B).

**Figure 5.**
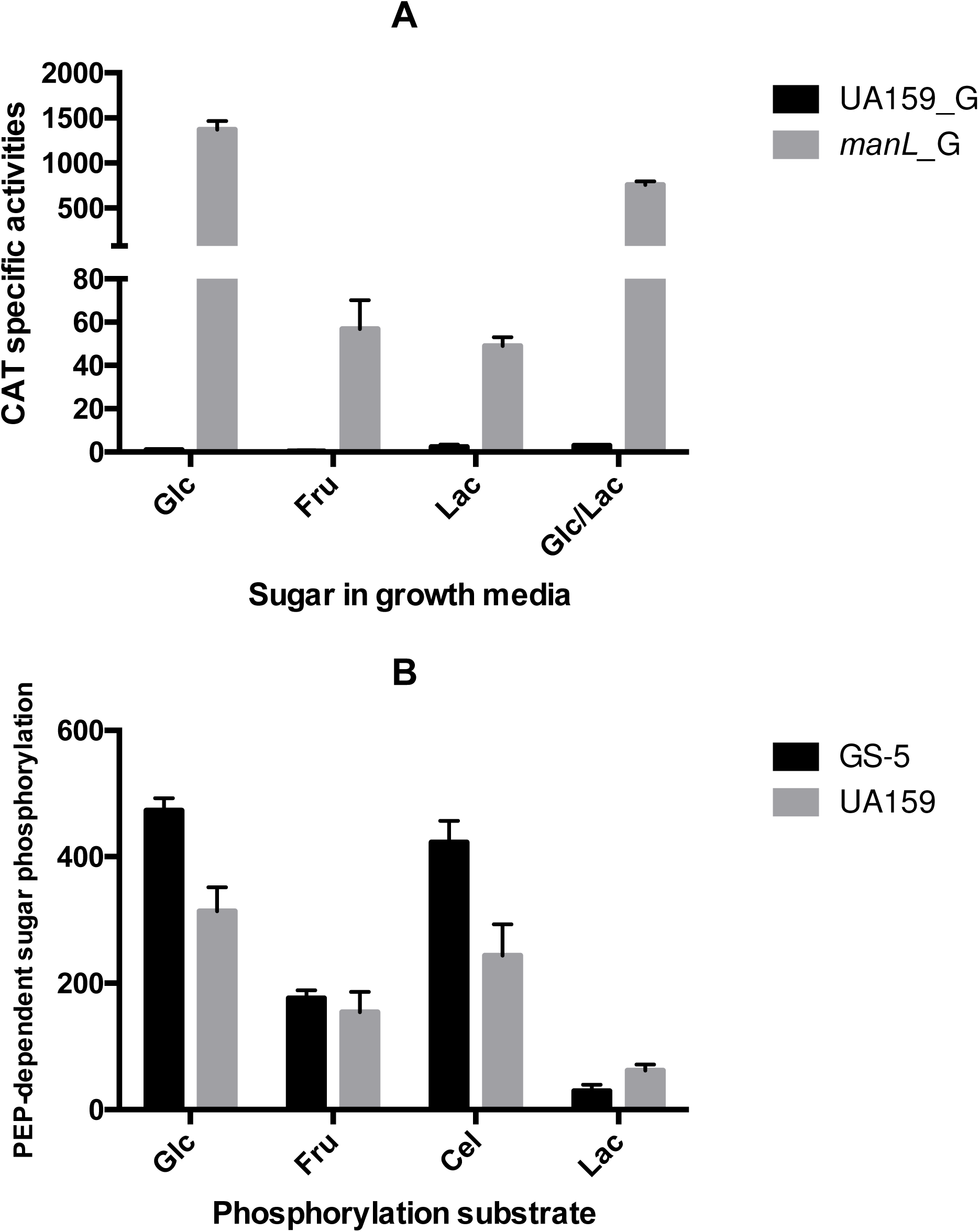
Expression of the cellobiose pathway by cells growing in the presence of lactose. (A) The activities of a P*celA::cat* reporter fusion were measured in wild-type UA159 or its *manL* mutant, with cells growing on glucose (Glc), fructose (Fru), lactose (Lac) or a mixture of 5 mM each of glucose and lactose. (B) PTS-dependent sugar phosphorylation assays were performed to measure the PTS activities transporting glucose, fructose, cellobiose (Cel) and lactose. Wild-type strains GS-5 and UA159 were cultured in FMC containing a mixture of 5 mM each of glucose and lactose before assay. All bars represent the averages from three independent experiments with each measurement repeated twice. The error bars represent the standard deviations.

In contrast, significantly more glucose was released by GS-5 than UA159 when growing on lactose (>1 mM/OD_600_) even for the wild type (Fig. 4). A *manL* mutant in the GS-5 genetic background accumulated more extracellular glucose (>3 mM/OD_600_) than the wild type. As re-internalization of glucose via EII^Man^ creates glucose-6-phosphate and other catabolites (e.g. F-1,6-bP) capable of triggering the activity of CcpA (2), the relaxed expression of the *lac* genes by the *manL* mutant of GS-5 is consistent with the conventional model for CCR. Therefore, it is plausible that the amount of glucose released during lactose metabolism is sufficient to trigger CcpA-dependent CCR in GS-5, as we observed in a previous study (6).

In light of the sequence divergence between the *lacA* promoters in GS-5 and UA159, the P*lacA::gfp* fusion created using the UA159 *lacA* promoter template, namely the version harboring only one (T)_7_G motif (Fig. S6), was introduced into the backgrounds of GS-5 and its *manL* mutant (Fig. 6). Interestingly, the *manL* mutant no longer showed enhanced expression of the reporter fusion when growing in the presence of lactose; instead it produced lower GFP signals than the wild type when cultured in a combination of glucose and lactose. In other words, the *lacA* expression patterns in these two strains now resembled those in the UA159 background. These results indicate that the difference in lactose metabolism observed between UA159 and GS-5 is partly attributable to how each *lac* operon is regulated by the glucose PTS. However, a brief analysis of the *lacA* promoter region in both strains failed to locate a conserved CcpA-binding site, but the potential that the (T)_7_G motif acts as cis-acting regulatory elements is supported by the data.

**Figure 6.**
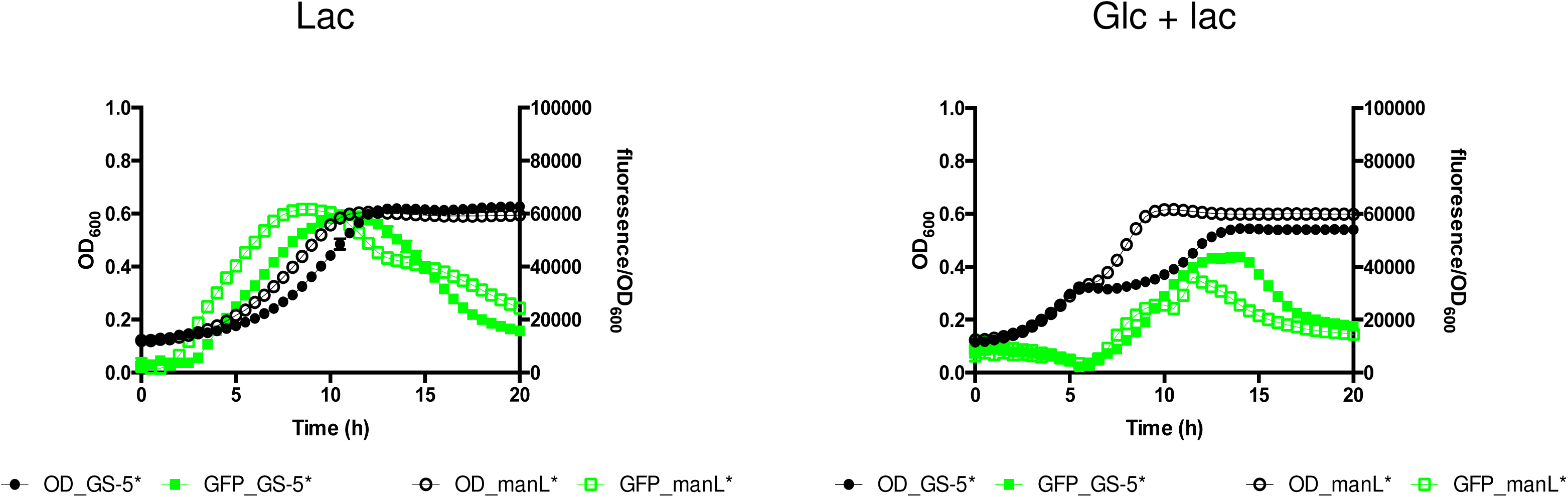
Expression of a UA159-derived P*lacA::gfp* fusion (relative fluorescence, green) in the GS-5 background. Wild type (filled circle) and a *manL* mutant (open circle) were cultivated to exponential phase in FMC containing 20 mM glucose, and then diluted into FMC containing 10 mM lactose (Lac), or 5 mM each of glucose and lactose (Glc + Lac). OD_600_ and GFP fluorescence were monitored using a BioTek Synergy 2 plate reader maintained at 37°C.

Lastly, when cellobiose or trehalose was used as the sole carbohydrate source to cultivate *S*. *mutans* deficient in EII^Man^, free glucose was again detected in the supernates, albeit at levels lower than what was released by lactose-grown cultures (Fig. 4 and data not shown). The cause for this difference has not been determined, but could reflect that glucokinase activity is increased when cells are cultured on disaccharides composed only of glucose.

### Free glucose and other preferred carbohydrates are required for efficient transition to growth on lactose

When exponential-phase cultures of *S*. *mutans* growing in FMC with 20 mM glucose or fructose were diluted 1:20 into fresh FMC containing lactose, carryover of residual glucose or fructose occurs, which could influence growth behaviors. To eliminate carryover of sugars and assess the impact of small amounts of hexose on the transition to lactose, cells were harvested in exponential phase and washed extensively with sterile phosphate-buffered saline (PBS), then small amounts (0, 0.125 mM, 0.25 mM) of glucose or other monosaccharides were added into the FMC-lactose medium before inoculation. Unexpectedly, without addition of the monosaccharides, washed cultures of *S. mutans* displayed an apparent growth arrest that lasted as long as 24 h (extended memory - Fig. 7, and Fig. S7 for more *S*. *mutans* isolates and commensals). Such growth arrest was particularly significant when the pre-cultures were prepared using FMC-fructose. However, addition of small amounts of the glucose, fructose, mannose, glucosamine (GlcN) or N-acetyl-glucosamine (GlcNAc) accelerated the transition of *S*. *mutans* to growth on lactose, drastically shortening the lag phase (Fig. 7). The effects of sugars on the duration of the lag phase were concentration-dependent and could be observed at levels of hexose as low as 10 μM (Fig. S8). Importantly, mutants lacking the PTS permease(s) for the cognate monosaccharides showed reduced or abolished responses to low levels of these hexoses (data not shown). When other disaccharides besides lactose were tested in this experiment, it was determined that similar effects on the lag phase for transition to growth on lactose could be observed for cellobiose or trehalose, but not for sucrose or maltose (data not shown). Certain *S*. *mutans* strains, however, were able to grow efficiently on some of these disaccharides alone, e.g., GS-5 on cellobiose. The fact that carbohydrates other than glucose were able to rescue growth on lactose suggests a more general effect by these preferred carbohydrates that impacts lactose metabolism, one that is likely independent of the cellobiose pathway discussed above.

**Figure 7.**
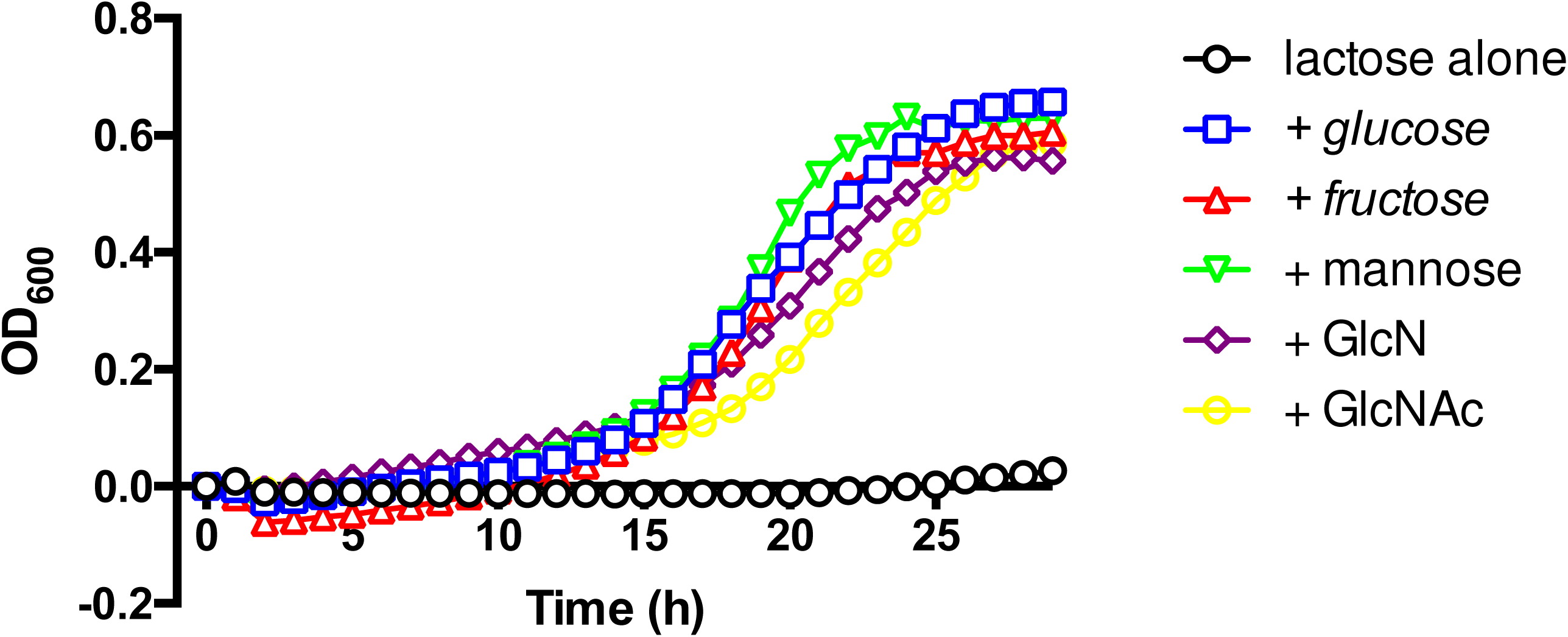
*S*. *mutans* requires preferred monosaccharides for growth on lactose. GS-5 was cultured to exponential phase in FMC that contained 20 mM glucose, washed three times with sterile PBS and re-suspended in the same buffer, then diluted 1:20 ratio into fresh FMC containing 10 mM lactose, or 10 mM lactose in addition to 0.125 mM of glucose, fructose, mannose, GlcN or GlcNAc, for growth monitoring in a Bioscreen C.

Also of particular interest was the fact that addition of galactose (as much as 0.25 mM) to any of the washed *S*. *mutans* cultures failed to rescue growth on lactose, even though a number of these isolates harbored genes encoding a high-affinity galactose-specific PTS permease (data not shown)(21). Further experimentation revealed that these isolates also showed a similar growth arrest when washed prior to transferring from glucose to galactose as the sole carbohydrate, a defect that again could be rescued by addition of small amounts of glucose (data not shown).

Collectively these results indicate the existence of distinct hierarchies in the preference for utilization of common carbohydrates by *S*. *mutans*, with most monosaccharides, sucrose and maltose being preferred, followed by trehalose, cellobiose, galactose and lactose. As *S*. *mutans* cultures have been shown to release and re-internalize glucose when growing on lactose, cellobiose or trehalose, often at concentrations that were comparable to what were used to elicit repression of genes for non-preferred carbohydrates, these findings suggest concentration-dependent and disparate effects of glucose and the glucose-PTS on catabolism of these less-preferred carbohydrates: the expression of the catabolic genes is negatively regulated by the glucose porter via CCR at higher concentrations of glucose, but efficient induction is dependent on small amounts of glucose, and apparently certain other sugars, for activation. The molecular mechanisms responsible for such activation remains to be elucidated, although it is worth noting that a positive feedback loop generally exists for the induction of these pathways (7, 9), where the signal output of these self-activating circuits could be enhanced not only by the substrates (for EII^Cel^) or metabolic intermediates (for EII^Lac^ and EII^Tre^), but also by triggering efficient expression of the PTS and the glycohydrolase required for catabolism of the non-preferred carbohydrate.

### *rel* mutants showed an altered ability to transition onto lactose

When faced with environmental stresses, such as amino acid starvation, bacteria often respond by synthesizing tetra- and penta-phosphorylated guanosine alarmones, collectively designated (p)ppGpp, that can suppress protein biosynthesis and alter gene regulation (22). There are three (p)ppGpp-metabolizing enzymes in *S*. *mutans*: RelA, which has both synthase and hydrolase activities, and two synthase-only enzymes, RelP and RelQ, which contribute to (p)ppGpp pools under specific circumstances (23). To examine the possibility that the stringent response is involved in the transition of *S*. *mutans* from one carbohydrate source to another, we studied the growth behaviors of a set of otherwise-isogenic mutants that lacked individual or combinations of the *rel* genes. As shown in Fig. 8 and Table 1, deletion of *relP* or *relQ* had no apparent impact on the ability of UA159 to transition from glucose onto lactose. On the other hand, deletion of the entirety of *relA* via allelic exchange (*relA*::Km), or the disruption of the (p)ppGpp-hydrolase function alone (*relA*:385)(24) resulted in an increase in the transition time for growth on lactose. Further, a strain missing all three *rel* enzymes (Δ*relAPQ*), which appears to produce no (p)ppGpp (23), also showed a similar increase in the time to transition to growth on lactose. Similar results were obtained from these mutants when the pre-cultures were prepared using FMC-fructose (data not shown). Previous work from our group characterizing the enzymatic activities of these mutants has indicated that while the triple-null mutant produces no (p)ppGpp, loss of the hydrolase activity of RelA (relA:385) or deletion of *relA* entirely results in higher basal levels of alarmones (24). Therefore, it appears that modest changes in (p)ppGpp pools resulting from loss of, or changes in the synthase/hydrolase capabilities of RelA, could significantly affect the ability of *S*. *mutans* to transition from a preferred carbohydrate onto a non-preferred source. It should be noted that our group has demonstrated that growth on fructose, as compared to glucose, induces changes in the transcriptome that alter the expression of many stress-related genes (13), and that accumulation of fructose-phosphate (14) and other sugar phosphates (25) could lead to growth arrest by *S*. *mutans*. Of note, *E*. *coli* has a requirement for an intact stringent response to cope with sugar-phosphate stress (26). We hypothesize that the PTS and presence of certain carbohydrate can contribute to prolonged growth arrest by altering (p)ppGpp pools, and that the effects can be influenced by the activity of PTS enzymes involved in the transport of preferred carbohydrates.

**Figure 8.**
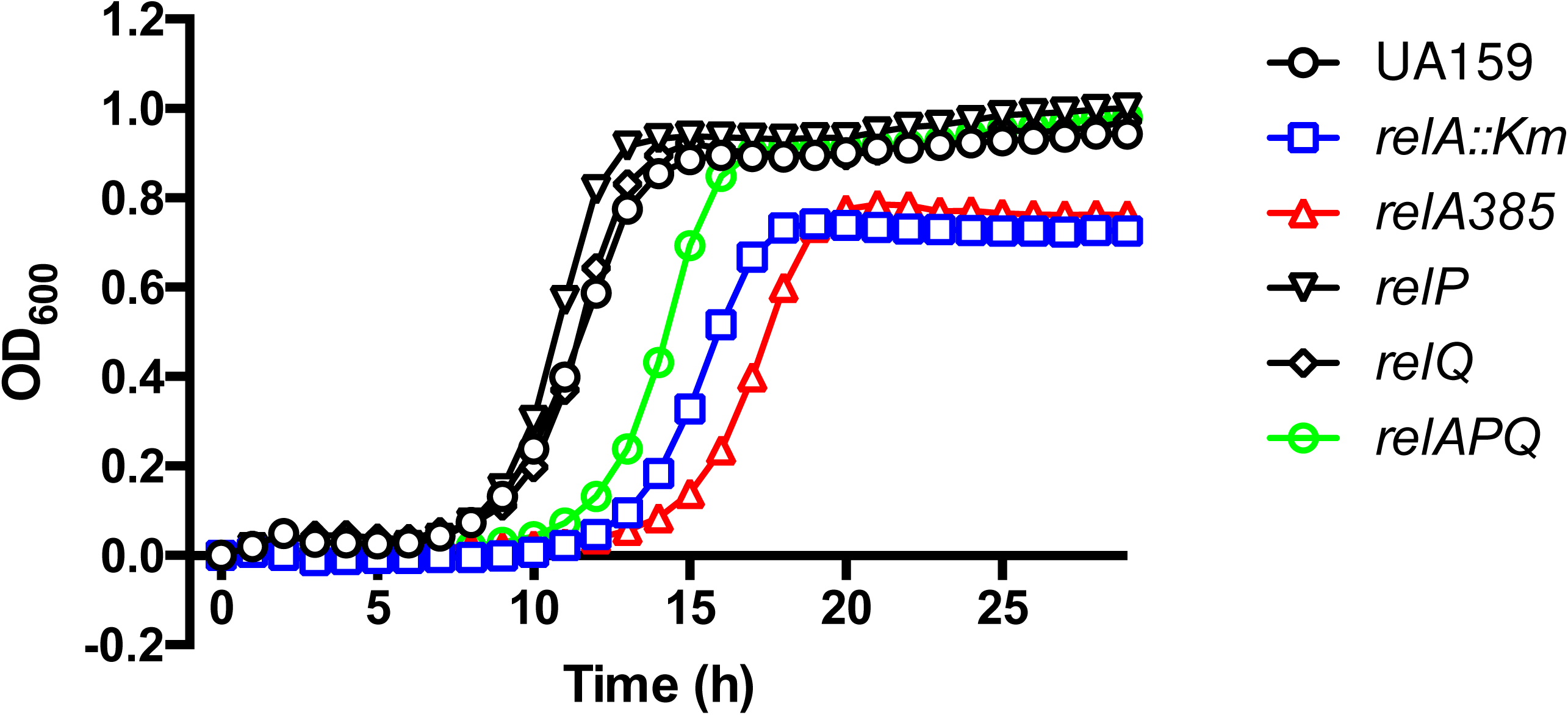
Mutations in (p)ppGpp-metabolizing enzymes affect growth of *S*. *mutans* on lactose after transferring from glucose. Wild-type strain UA159 and its isogenic mutants with complete deletion of *relA (relA::Km)*, *relP*, *relQ*, all three (*relAPQ*), or a truncation in *relA* (relA385) that abolishes its (p)ppGpp-hydrolase activities, were each cultured in FMC containing 20 mM glucose to mid-exponential phase, then diluted 1:20 into fresh FMC constituted with 10 mM lactose for growth monitoring.

### Summary and relevance to oral microbial ecology

Biofilms in the human oral cavity experience frequent and substantial shifts in the availability and type of carbohydrates, and most bacteria depend on CCR to prioritize sugar metabolism. CCR in *S*. *mutans* relies heavily on the PTS for CCR and micromolar levels of preferred carbohydrates can trigger PTS-dependent CCR for genes for secondary metabolites, such as *fruA* encoding a fructan hydrolase (6). Here, we investigate the unusual phenomenon of long-term memory and explored the molecular basis for the delay in adaptation by *S*. *mutans* to the introduction of lactose depending upon prior exposure to glucose or fructose. We also show that *S*. *mutans* requires low levels of preferred carbohydrates (including glucose) for efficient induction of the *lac* operon, notable also because substantial amounts of free glucose are released into the environment by cells that are actively catabolizing lactose (as well as cellobiose and trehalose). Therefore, as a byproduct of metabolism of lactose and other disaccharides, free glucose at low concentrations may serve as a signal to prime the induction of the *lac* operon, however higher levels of glucose or other preferred carbohydrates could down-regulate *lac* expression through CCR. This phenomenon echoes a recent report wherein glucose-PTS of *Streptococcus pneumoniae* was shown to serve as a surveyor of external carbohydrates that is required for the metabolism of multiple carbohydrates (27). Unlike for the pneumococcus, fructose, which is frequently present in the human diets, but is also a byproduct of sucrose metabolism by extracellular enzymes and the PTS of *S*. *mutans* (12), could be viewed as a negative signal for the induction of the *lac* system. We posit that, as a bet-hedging strategy, certain populations of *S*. *mutans* respond to a combination of these hexoses in a way that delays the expression of the *lac* operon in anticipation of future ingestion of fructose or sucrose by the host, in order to avoid the energy expenditure required for synthesizing enzymes for intermittently available, less-preferred carbohydrates. We have previously shown that growth on fructose or sucrose down-regulates the expression of enzymes required for glycogen metabolism (SMU.1535 ~ SMU.1539) (13), whereas a FruK (1-phosphofructokinase)-deficient mutant overexpresses some of the same enzymes (SMU.1536 ~ SMU.1539), presumably in response to increased levels of fructose-1-phosphate (14). It remains to be clarified how these changes in glycogen-related gene expression affect the persistence of the bacterium under these conditions. Finally, this study raises the question of how free hexoses released during metabolism of these disaccharides influence oral biofilm ecology, since other microorganisms in the same niche can be provided with preferred carbohydrates when *S*. *mutans* and possibly other oral streptococci metabolize disaccharides. One scenario of particular interest to us is the emergence of so-called cheaters within the population of *S*. *mutans* that take advantage of these free hexoses without investing in the production of the enzymes necessary for catabolizing lactose or other disaccharides, as indicated by our preliminary study (Fig. 5B and data not shown). Further research at the single-cell level could help to shed more light on the behavior of individual cells under these conditions.

## MATERIALS AND METHODS

### Bacterial strains, media and culture conditions

*Streptococcus mutans*, including wild type and mutant strains, plus five other lactic acid bacteria (LAB, Table 2) were maintained on brain heart infusion (BHI)(Difco Laboratories, Detroit, MI) agar plates or in liquid medium at 37°C in a 5% CO_2_, aerobic atmosphere. Antibiotics were added to the media, when necessary: kanamycin (0.5 ~ 1 mg ml^-1^), erythromycin (5 ~ 10 μg ml^-1^), spectinomycin (0.5 ~ 1 mg ml^-1^), and tetracycline (10 μg ml^-1^). *Escherichia coli* DH10B was used as a cloning host and cultivated in L-broth at 37°C with agitation in air. When needed, L-broth was supplemented with antibiotics: kanamycin (50 μg ml^-1^), and erythromycin (200 μg ml^-1^). For most experiments, *S*. *mutans* and other LAB were cultured in a modified version of the chemically defined medium FMC (28) formulated with various carbohydrates (Thermo Fisher Scientific, Waltham, MA; and MilliporeSigma, St. Louis, MO), as indicated.

**Table 2.** Bacterial strains used in this study.

| Strains | Relevant characteristics<sup>a</sup> | Source or reference |
| --- | --- | --- |
| ***Streptococcus mutans*** |  |  |
| UA159 | Wild type, serotype *c* | University of Alabama |
| JAM1 | UA159 *manL::Km* | (35) |
| MMZ540 | UA159 *celR::Sp manL::Em* | (7) |
| JLrelA | UA159 *relA::Km* | (36) |
| *relA::385* | UA159 containing a truncation of *relA* starting at 385th residue | (24) |
| JLrelP | UA159 *relP::Km* | (23) |
| JLrelQ | UA159 *relQ::Km* | (23) |
| JLT1 (Δ*relAPQ*) | UA159 *relA::Em relP::Sp relQ::Km* | (23) |
| MMZ712 | UA159 *lacE::Em* | (9) |
| MMZ702 | UA159 *lacG::Em* | (9) |
| MMZ1285 | UA159 *PlacA::gfp* | This study |
| MMZ1287 | UA159 *manLMN::Em PlacA::gfp* | This study |
| MMZ1548 | UA159 *manL::Em PlacA::gfp* | This study |
| MMZ493 | UA159 *PcelA::cat* | (7) |
| MMZ495 | UA159 *manL::Em PcelA::cat* | (7) |
| MMZ1422 | UA159 *glk::Km* | This study |
| MMZ1455 | UA159 *glk::Km manL::Em* | This study |
| GS-5 | Wild type, serotype *c* | R. J. Gibbons |
| MMZ1462 | GS-5 *manL::Em* | This study |
| MMZ1424 | GS-5 *celR::Km* | This study |
| MMZ1509 | GS-5 *celA::Em* | This study |
| MMZ1504 | GS-5 *celB::Em* | This study |
| MMZ1510 | GS-5 *lacE::Em* | This study |
| MMZ1517 | GS-5 *lacG::Em* | This study |
| MMZ1480 | GS-5 *PlacA::gfp* | This study |
| MMZ1483 | GS-5 *manL::Em PlacA::gfp* | This study |
| LM-7 | Wild type, serotype *e* | A. Eisenberg |
| V403 | Wild type, serotype *c* | T. Kitten |
| 10449 | Wild type, serotype *c* | A. Eisenberg |
| Smu44 (11VS1) | Wild type, serotype *e* | (37) |
| Smu33 (11SSST2) | Wild type, serotype *c*, EII<sup>Gal</sup> + | (37) |
| Smu21 (1SM1) | Wild type, serotype *e*, EII<sup>Gal</sup> + | (37) |
| UA101 | Wild type, serotype *c* | Laboratory strain |
| OMZ175 | Wild type, serotype *f*, EII<sup>Gal</sup> + | B. Guggenheim |
| **Others** |  |  |
| *Streptococcus* |  |  |
| *gordonii* DL1 | Wild type standard strain | ATCC 49818 |
| *Streptococcus* |  |  |
| *sanguinis* SK150 | Wild type standard strain | Laboratory stock |
| *Streptococcus* |  |  |
| *intermedius* | Wild type standard strain | ATCC 31412 |
| *Streptococcus* |  |  |
| *salivarius* 57.I | Wild type standard strain | (38) |
| *Lactococcus lactis* | Wild type standard strain | ATCC 11454 |

<sup>a</sup> *Km* indicates kanamycin resistance; *Sp*, spectinomycin; *Em*, erythromycin.

To monitor bacterial growth, all strains were cultivated overnight in 5% CO_2_ in FMC containing the specified carbohydrate, diluted 1:10 into the same medium and cultured until the optical density at 600 nm (OD_600_) reached 0.5. Then, the cultures were again diluted (1:20) into fresh FMC constituted with the desired carbohydrates and incubated at 37°C in a Bioscreen C Lab system (Helsinki, Finland). Each experiment included three individual isolates (biological replicates) in triplicate (technical replicates), and each sample (300 μl) was overlaid with sterile mineral oil (60 μl) to reduce exposure to air, and the OD_600_ was measured every hour after a brief agitation. In experiments designed to eliminate carryover of sugar(s) from the prior incubation when cultures were diluted, exponential-phase cultures were first cooled on ice, harvested by centrifugation, and then washed 3 times with an equal volume of cold, sterile PBS. The cultures were then resuspended in the desired medium, diluted as necessary, and growth was monitored as above.

For bacteria containing a P*lacA::gfp* fusion, cells were cultured as above to exponential phase, diluted 1:20 into 200 μl FMC medium in a Costar (Corning, Corning, NY) 96-well microtiter plate with clear bottom, and the culture was overlaid with 60 μl of mineral oil. The growth of the bacteria (OD_600_) and expression of the promoter fusion as represented by green fluorescent light emission (excitation 485/20 nm, emission 528/20 nm) were monitored simultaneously every 30 min for 20 h using a fluorescence-capable Synergy 2 Multi-Mode reader from BioTek (Winooski, VT). For each test strain, a control strain for background subtraction was included; the control had the same genetic background, but lacked the reporter fusion. To calculate the relative expression of the promoter fusion, the background fluorescence of the control sample was subtracted from each raw reading of the test sample and then divided by the corresponding OD_600_ value for normalization. Finally, cultures used for measurement of glucose in culture supernates, and for CAT and PTS assays, were also prepared by diluting from an overnight culture using fresh FMC containing the specified carbohydrate(s), followed by incubation until OD_600_ reached 0.5.

### Construction of genetic mutants and promoter::*gfp* fusions

Standard recombinant DNA techniques were employed to engineer plasmids and mutant strains. All restriction enzymes were purchased from New England Biolabs (Beverly, MA) and used as directed by the supplier. DNA preparation was performed using Qiaquick DNA purification kits supplied by Qiagen (Valencia, CA). All DNA oligonucleotides were synthesized by Integrated DNA Technologies (Coralville, IA) and listed in Table 3.

**Table 3.**
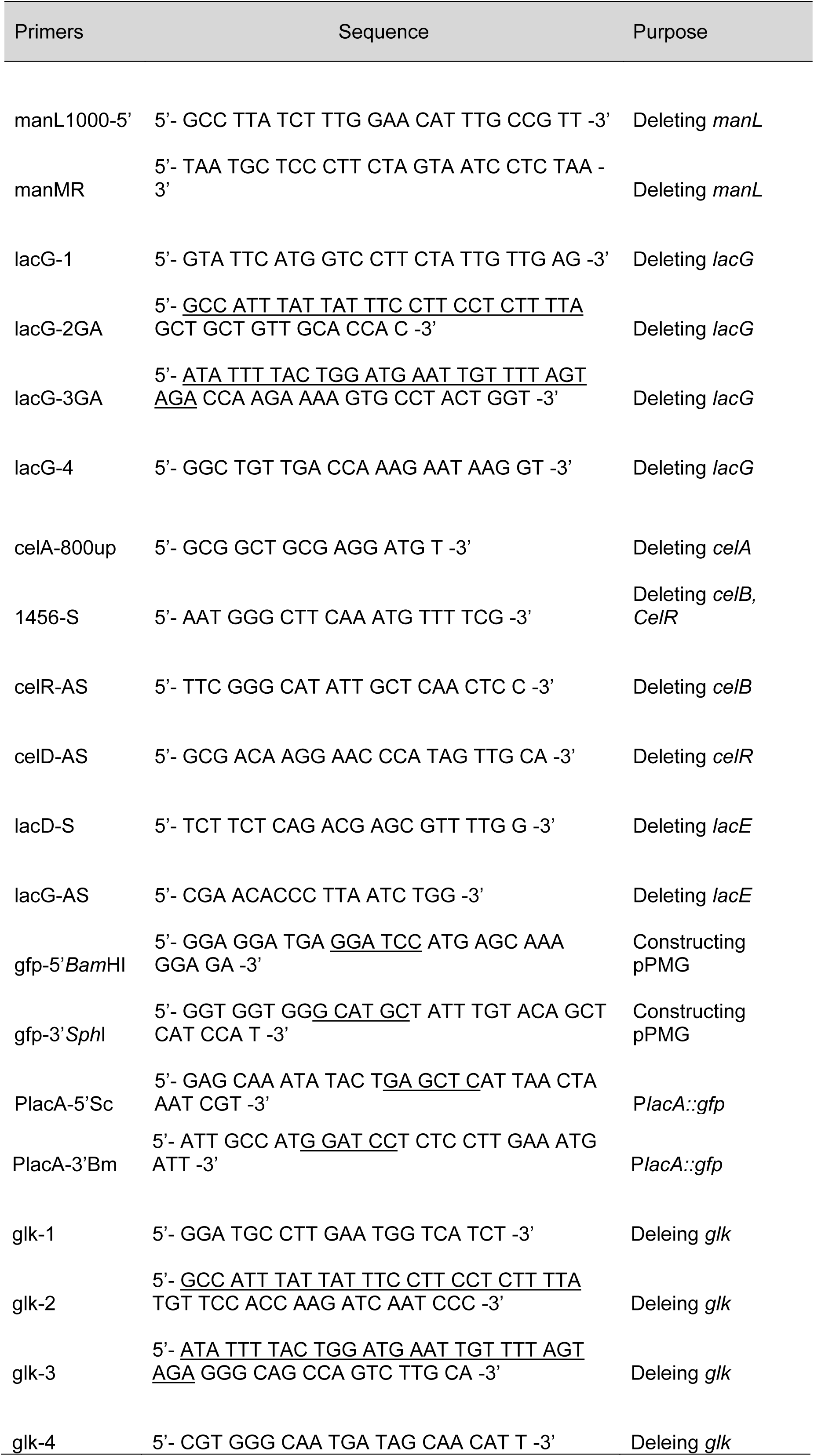
Primers used in this study.

The majority of the mutant derivatives of *S*. *mutans* used in this study were engineered by following the PCR-ligation approach (29) utilizing three genetic cassettes encoding for resistance to kanamycin (*Km*), erythromycin (*Em*) or spectinomycin (*Sp*). To construct allelic exchange mutants of *lacG* and *glk*, however, two PCR products flanking each target gene were created so that sufficient sequence overlap (>25 bp) existed between these DNA fragments and the antibiotic cassette (see Table 3 for detail). A Gibson Assembly mastermix (New England Biolabs) was then utilized to ligate these DNA fragments in a one-step, 1-h reaction before transformation of *S*. *mutans*. Based on the high degrees of sequence similarity between UA159 and GS-5, a number of mutants in GS-5 background were constructed by transformation of GS-5 using PCR products derived from the UA159 mutants, which contained the flanking sequences and an antibiotic marker in place of the target gene.

To construct the P*lacA::gfp* reporter fusion, a plasmid pPMG containing a superfolder GFP protein (sGFP)(30) was engineered on the basis of a *cat*-fusion plasmid pJL84, for the purpose of integrating exogenous DNA at the *phnA*-*mtlA* locus (31). A set of primers, gfp-5’BamHI and gfp-3’SphI (Table 3), were designed for the amplification of the *gfp* gene, using plasmid pCM11 (32) as the template. After digestion with restriction enzymes *BamHI* and SphI, the *gfp* fragment was cloned into pJL84 that was treated with the same enzymes to drop out the *cat* gene, resulting in plasmid pPMG. Subsequently, the *lacA* promoters from UA159 and GS-5 were amplified using primers PlacA-5Sc and PlacA-3Bm (Table 3) and directionally cloned into *Sac*I and *BamH*I sites in pPMG. These plasmids were then used to transform various *S*. *mutans* strains for integration of the reporter fusions. All plasmids and genetic mutants engineered in this study were verified by sequencing or PCR followed by sequencing, respectively, in order to exclude any clones with unintended mutations in the target genes and in genomic regions immediately adjacent to them.

### Biochemical assays

Glucose in the supernates of bacterial cultures were measured using a glucose oxidase kit and a hexokinase kit (MilliporeSigma), following protocols provided by the supplier. Chloramphenicol acetyltransferase activity (CAT)(33) and PEP-dependent sugar phosphorylation (PTS)(34) assays were performed according to protocols established in our laboratory, as detailed elsewhere (31).

## ACKNOWLEDGEMENTS

We appreciate the technical support from Lulu Chen in performing some of the growth assays. This study was supported by DE12236 from the National Institute for Dental and Craniofacial Research.

